# Unique maternal immune and functional microbial profiles during prenatal stress

**DOI:** 10.1101/2020.05.26.116574

**Authors:** Adrienne M. Antonson, Morgan V. Evans, Jeffrey D. Galley, Helen J. Chen, Therese A. Rajesekera, Sydney M. Lammers, Vanessa L. Hale, Michael T. Bailey, Tamar L. Gur

**Author notes:** Corresponding author information*: Tamar L. Gur, MD PhD, 120A Institute for Behavioral Medicine Research Building, 460 Medical Center Drive, Columbus OH, 43210.

## Abstract

Maternal stress during pregnancy is widespread and stress-induced fetal neuroinflammation is thought to derive from a disruption in intrauterine immune homeostasis, though the exact origins are incompletely defined. We aimed to identify divergent immune and microbial metagenome profiles of stressed gestating mice that may underlie detrimental inflammatory signaling at the maternal-fetal interface. In response to stress, maternal glucocorticoid circuit activation corresponded with diminished spleen mass and IL-1β production, reflecting systemic immunosuppression. At the maternal-fetal interface, density of placental mononuclear leukocytes decreased with stress. Yet maternal whole blood leukocyte analysis indicated monocytosis and classical M1 phenotypic shifts. Genome-resolved microbial metagenomic analyses revealed reductions in genes, microbial strains, and metabolic pathways in stressed dams that are primarily associated with pro-inflammatory function. Overall, these data indicate that stress disrupts maternal immunological and microbial regulation during pregnancy, characterized by concurrent anti- and pro-inflammatory signatures, which may displace immune equilibrium at the maternal-fetal interface.

## 1. Introduction

Prenatal psychological stress, or maternal stress during pregnancy, has been linked to poor birth outcomes and offspring mental health disorders. While the mechanisms underlying fetal programming during prenatal stress are numerous and incompletely defined^1^, long-term adverse mental and physical outcomes in offspring are well documented^2^. The incidence of prenatal stress, which is not confined to diagnosable mental health disorders such as anxiety and depression but also includes taxing major life events, daily hardships, and exposure to disaster^2^, is underestimated and underreported. Yet there is evidence that the majority of pregnant women experience some form of stress during pregnancy, with estimates as high as 84 percent^3^. Variations in maternal physiology that occur during stress, including activation of the hypothalamic-pituitary-adrenal (HPA)-axis, altered immune profiles, and disrupted microbial homeostasis are thought to play a major role in determining offspring outcomes.

Investigations into the unique immune profile of pregnancy, apparent within tissues of the maternal-fetal interface^4^ and in maternal circulation^5^, have revealed that immune tolerance of fetal antigen throughout gestation is much more complex than a physical barrier between maternal and fetal tissues^6^. Communication between placental trophoblast cells and uterine immune and endometrial cells provides an intricate regulation of the inflammatory microenvironment throughout pregnancy^4^. The majority of pregnancy is characterized as anti-inflammatory and immunosuppressive, with maternal leukocyte phenotypes generally reflecting an alternatively activated T_H_2- and M2-like profile in order to support symbiotic fetal growth and development. However, pro-inflammatory milieus bookend this extended period and regulate the early stage of blastocyst implantation and placentation, as well as the late stage of parturition and labor^4^. Therefore, responsive maternal immune adaptations that coincide with these developmental stages are necessary for maintaining a healthy pregnancy^7^. Altered proportions of these otherwise well-controlled leukocyte populations are associated with pregnancy complications like preterm birth^8^, preeclampsia^9^, and miscarriage^10,11^.

While circulating maternal leukocyte populations change in response to gestation progression^12^, healthy pregnancy is predominantly characterized by monocytosis and neutrophilia^13^. Pregnancy-specific increases in a unique subset of immunosuppressive cells of myeloid origin has also recently received attention. These myeloid-derived suppressor cells (MDSCs), which can further be classified as polymorphonuclear (PMN) or monocytic (M), are an integral part of maintaining an immune balance at the maternal-fetal interface^6^. In mice, PMN-MDSCs bear many of the same markers as neutrophils (CD11b^+^Ly6G^+^Ly6C^low^SSC^hi^), while M-MDSCs resemble inflammatory monocytes (CD11b^+^Ly6G^-^Ly6C^hi^SSC^low^)^10^. While the potential disruption of gestational monocytosis or neutrophilia by psychological stress has not been fully explored, stress-activated signaling pathways have been proposed as possible triggers leading to altered immune profiles in pregnancy disorders^14^. Indeed, stress stimulates hematopoiesis and increases the release of neutrophils and monocytes from the bone marrow into circulation^15^. Concomitant activation of the HPA-axis and release of glucocorticoids results in a redistribution of these leukocytes from circulation into various tissues^16^, potentially including the uterus and placenta. Increases in circulating glucocorticoids can also act directly on monocytes to augment expression of chemokine receptor CCR2^17^, enhancing cell migratory capacity through CCL2-CCR2 mediated chemotaxis. In pregnancy, CCL2-mediated recruitment of leukocytes (including MDSCs) to the maternal-fetal interface has been demonstrated, and is dependent upon production and release of these chemokines by placental trophoblast cells^4,18^. This orchestrated process of leukocyte recruitment to decidual and placental tissue is necessary for successful pregnancy^4^ and altered production of chemokines like CCL2 have been linked to poor pregnancy outcomes^19^. The specific effects of prenatal stress on recruitment of leukocytes to the maternal-fetal interface, however, has not been explored.

Altered immune responses occurring in cases of prenatal stress^20^ or pregnancy complications^11^ may also be related to disrupted microbial balance^8,21^, even in the absence of an overt infection. Animal models of psychological stress demonstrate a shift in gastrointestinal (GI) microbiota composition and function^22–24^, and there is evidence that this is also true during pregnancy^25,26^. Shifts in microbial homeostasis during gestation can impact establishment of the founding infant microbiome^27–30^ and have long-term implications for offspring immune development^31,32^. Whether gestational insults also shift endogenous microbes at reproductive tissue sites is still controversial^33,34^, although translocation of microbes and bacterial peptidoglycan across the maternal-fetal interface during pregnancy has been documented in rodents^35,36^. Stressor-induced disruptions in GI microbe populations can compromise intestinal barrier integrity, leading to release of microbes into circulation and subsequent microbicidal activation of splenic macrophages^37^. The structure of the microbiome influences the reactivity of the immune and inflammatory responses, and dysbiotic microbiome profiles, particularly those associated with GI metabolism, are often thought to precede disease development^38–41^. While there is a potential for bacterial signaling to potentiate leukocyte expansion, trafficking, and pro-inflammatory activation at the maternal-fetal interface, this has not been examined during prenatal stress.

Using a murine model of psychological restraint stress during gestation, we tested the hypothesis that stress activates the HPA-axis and disrupts the endogenous GI microbial community, augmenting leukocytosis and subsequent leukocyte trafficking to uterine and placental tissues. Previously, we have shown that our model of prenatal stress leads to increased inflammation in the placenta and fetal brain, and results in aberrant behaviors and neuroinflammation in adult offspring^25,42^. We recently demonstrated that stressor-induced fetal brain and placental inflammation is largely absent in CCL2 knock-out (KO) and germ-free animals, and that CCL2 KO adult offspring are protected from deficits in sociability^43^. While our data indicate that CCL2-signaling and endogenous microbes play integral roles in mediating prenatal stress outcomes^43^, it is not yet known whether these factors may also coincide with an infiltration of immune cells to the maternal-fetal interface. Here, we examined maternal endocrine and immune outcomes in order to establish a comprehensive gestational stress phenotype and to determine whether leukocyte trafficking to the maternal-fetal interface augments detrimental intrauterine inflammation.

## 2. Results

### 2.1. Psychological stress restricts maternal body weight gain trajectory during gestation

Pregnant dams were subjected to repeated restraint stress (2 h/day) from gestational day (GD)10 to GD16, and tissues were collected on GD17 (**Fig. 1A**). As expected, dam body weight gained across gestation (GD0 to GD16) correlated with litter size for both treatment groups (**Fig. 1B**). However, linear regression analyses revealed a restricted body weight gain trajectory in stressed dams (NS vs. S y-intercept comparison, p < 0.05; **Fig. 1B**), resulting in less weight gained per pup when maternal weight gain was normalized to litter size (**Fig. 1C**). All dams started at the same pre-breeding body weight regardless of treatment, and, when not corrected for litter size, body weight gained across the first 10 and 16 days of gestation did not differ (**Supp. Table 1**). Litter size and total number of fetal resorptions were not impacted by stress (**Supp. Table 1**). Together, these data indicate that repeated restraint stress during mid-to-late gestation limits normal pregnancy weight gain trajectories without compromising overall litter size or fetus viability.

**Fig. 1.**
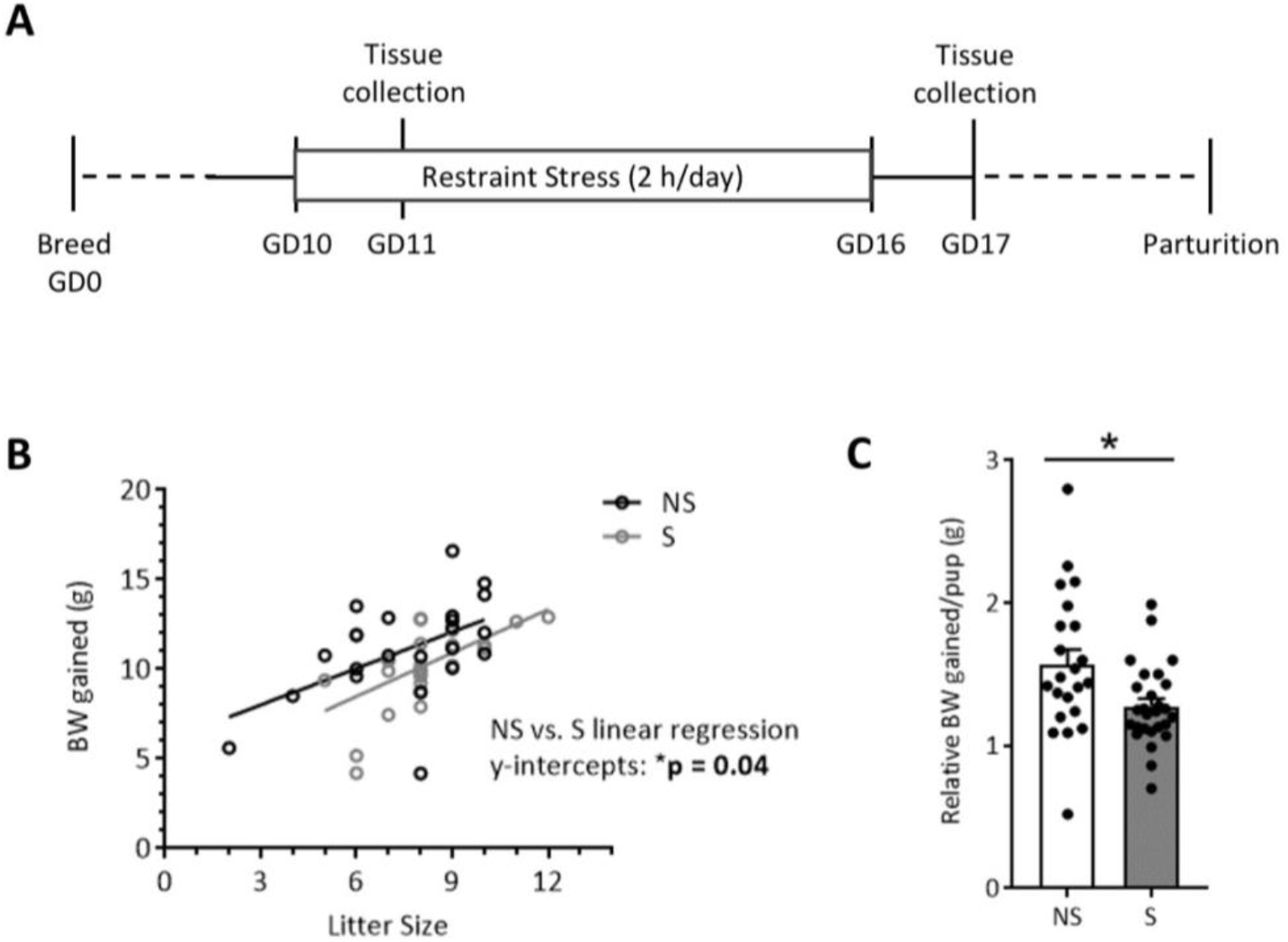
Restraint stress restricts gestational weight gain. The study design is depicted in **(A)**. While body weight gain **(B)** positively correlated with litter size regardless of treatment (NS: R^2^= 0.27, p = 0.01; S: R^2^= 0.34, p = 0.001; linear regression slope comparison, p = 0.71), linear regression y-intercepts differed across treatment groups (p = 0.04), reflecting a reduced weight gain from GD0 to GD16 with stress. This is further reflected when weight gain is normalized to litter size, revealing a **(C)** decreased weight gain per pup in the stressed group. Data are mean ± SEM. NS = non-stressed; n = 22, S = stressed; n = 27. * = p < 0.05.

### 2.2. Maternal glucocorticoid and systemic immune circuits are shifted in response to stress

Restraint stress during gestation raised circulating serum corticosterone levels (**Fig. 2A**) and increased relative gene expression of corticosteroid releasing hormone (*CRH*) in the hypothalamus and tended to increase glucocorticoid receptor (*NR3C1*) in the amygdala (p = 0.099; **Fig. 2B**). The above changes were concomitant with systemic indicators of immunosuppression, reflected by a decrease in spleen weight and in relative concentrations of splenic IL-1β, but not TNFα (**Fig. 2C**). Splenic gene expression is presented in **Supp. Table 2** and revealed no effect of stress upon immune genes. Circulating serum levels of inflammatory cytokines, measured through multiplex assay, were also relatively unchanged (**Supp. Table 3**) or below quantitative range (IFN-γ, IL-2, IL-4, and IL-12p70).

**Fig. 2.**
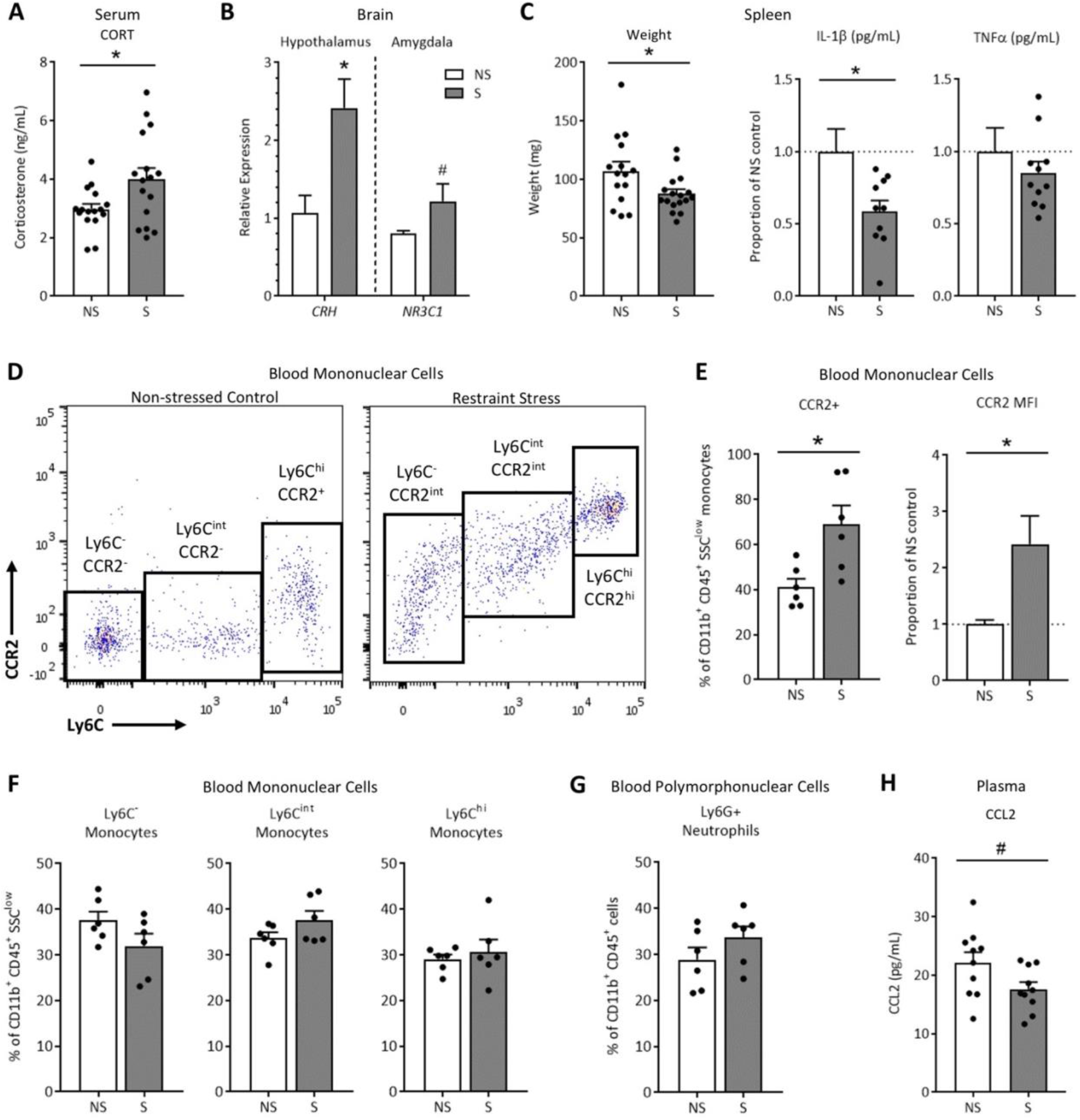
Restraint stress impacts systemic glucocorticoid and immune circuits. Stressing pregnant dams **(A)** increased serum corticosterone and **(B)** upregulated expression of *CRH* in the hypothalamus and glucocorticoid receptor (*NR3C1*) in the amygdala (p = 0.099). Immunosuppression was indicated by reduced **(C)** spleen weights and relative concentration of splenic IL-1β, but not TNFα (p = 0.37). A general increase in CCR2 expression is demonstrated in **(D)** representative scatter plots of whole blood CD11b^+^CD45^+^Ly6G^-^SSC^low^ mononuclear cells (Mo/M-MDSCs). When quantified, CCR2 expression increased within the mononuclear cell population as a whole, evidenced by **(E)** percent CCR2^+^ cells and CCR2 median fluorescence intensity (MFI). However, populations of mononuclear **(F)** Ly6C^-^CCR2^-^ alternative M2 monocytes (p = 0.12), Ly6C^int^CCR2^-^ transitional monocytes (p = 0.14), and Ly6C^hi^ CCR2^+^ classical M1 monocytes (p = 0.59) did not differ, nor were there overt differences in populations of circulating **(G)** CD11b^+^CD45^+^Ly6G^+^ polymorphonuclear cells (neutrophils or PMN-MDSCs; p = 0.21). Circulating plasma chemokine CCL2 concentrations **(H)** tended to decrease with stress (p = 0.056). Data are mean ± SEM, with each dot representing a single dam. NS = non-stressed, S = stressed; Mo/M-MDSCs = monocyte/monocytic myeloid-derived suppressor cells. CORT assay: n = 16/group; Spleen mass: n = 21-26/group; Spleen protein: n = 6-11/group; Gene expression: n = 7-10/group; Flow cytometry: n = 6/group. * = p < 0.05, # = p < 0.10.

As stress has been shown to recruit monocytes from the bone marrow into circulation (i.e. monocytosis) in part through CCL2-CCR2 chemotaxis, whole blood leukocytes were examined using flow cytometry. Cells were gated on double-positive expression of CD11b and CD45, and then further gated based on SSC properties. Examination of mononuclear cells (i.e. monocytes or M-MDSCs) revealed a notable shift towards CCR2^hi^ in stressed pregnant dams (scatter plots depicted in **Fig. 2D**). Quantification of this shift revealed that the percent of CD11b^+^CD45^+^SSC^low^ cells positive for CCR2 (encompassing low, intermediate, and high expression levels) increased, and overall median fluorescence intensity of CCR2 within these cells more than doubled (**Fig. 2E**). However, the percent of CD11b^+^CD45^+^SSC^low^ cells displaying each of the three monocyte phenotypes did not significantly differ (**Fig. 2F**), nor did overall leukocyte counts (**Table 1**). Ly6G^+^ polymorphonuclear cells (i.e. neutrophils or PMN-MDSCs) also did not overtly shift (**Fig. 2G**). Interestingly, circulating levels of plasma CCL2 tended to *decrease* with stress by GD17 (p = 0.056; **Fig. 2H**). These data indicate that repeated restraint stress activates glucocorticoid circuits, likely leading to systemic immunosuppression. Concomitant increases in CCR2 within circulating monocytes suggest a possible recruitment of CCR2^+^ mononuclear cells from the bone marrow and/or classical M1 activation, despite a trending reduction in plasma CCL2 at this time point.

**Table 1.** Circulating leukocyte counts.

| Leukocyte type (CD45<sup>+</sup>CD11b<sup>+</sup>) | Non-stressed | Stressed | p-value |
| --- | --- | --- | --- |
| Ly6G<sup>+</sup> (neutrophils) | 467.4 ± 53.1 | 487.1 ± 31.1 | 0.75 |
| SSC<sup>low</sup> (total monocytes) | 761.2 ± 90.2 | 721.5 ± 75.4 | 0.75 |
| SSC<sup>low</sup> Ly6C<sup>-</sup> (alternative monocytes) | 303.9 ± 53.0 | 229.4 ± 42.5 | 0.30 |
| SSC<sup>low</sup> Ly6C<sup>int</sup> (intermediate monocytes) | 247.7 ± 20.4 | 271.9 ± 21.2 | 0.45 |
| SSC<sup>low</sup> Ly6C<sup>hi</sup> (classical monocytes) | 209.7 ± 19.8 | 221.6 ± 31.8 | 0.77 |

Circulating leukocyte counts (cells/μL) from non-stressed and stressed gestating dams. All cells are CD45<sup>+</sup>CD11b<sup>+</sup>. Unpaired T tests; data are presented as mean ± SEM; n = 6/group.

### 2.3. The uterine immune environment at GD17 appears resistant to psychological stress

Investigation of uterine CD11b^+^CD45^+^ leukocyte populations revealed that this reproductive tissue displays no overt changes in percent of polymorphonuclear cells (neutrophils or PMN-MDSCs; **Fig. 3A**) or mononuclear cells (monocytes [Mo], macrophages [Mϕ], and M-MDSCs; **Fig. 3B**). Unlike whole blood, median fluorescence intensity of CCR2 within the Mo/Mϕ/M-MDSCs population did not differ from controls (**Fig. 3C**). Expression of immune genes within uterine tissue, including chemokine *CCL2*, remained stable (**Supp. Table 4**), and relative CCL2 protein concentrations did not differ (data not shown; p = 0.62; proportion of NS control: NS: 0.99 ± 0.25, n = 12; S: 0.82 ± 0.19, n = 7).

**Fig. 3.**
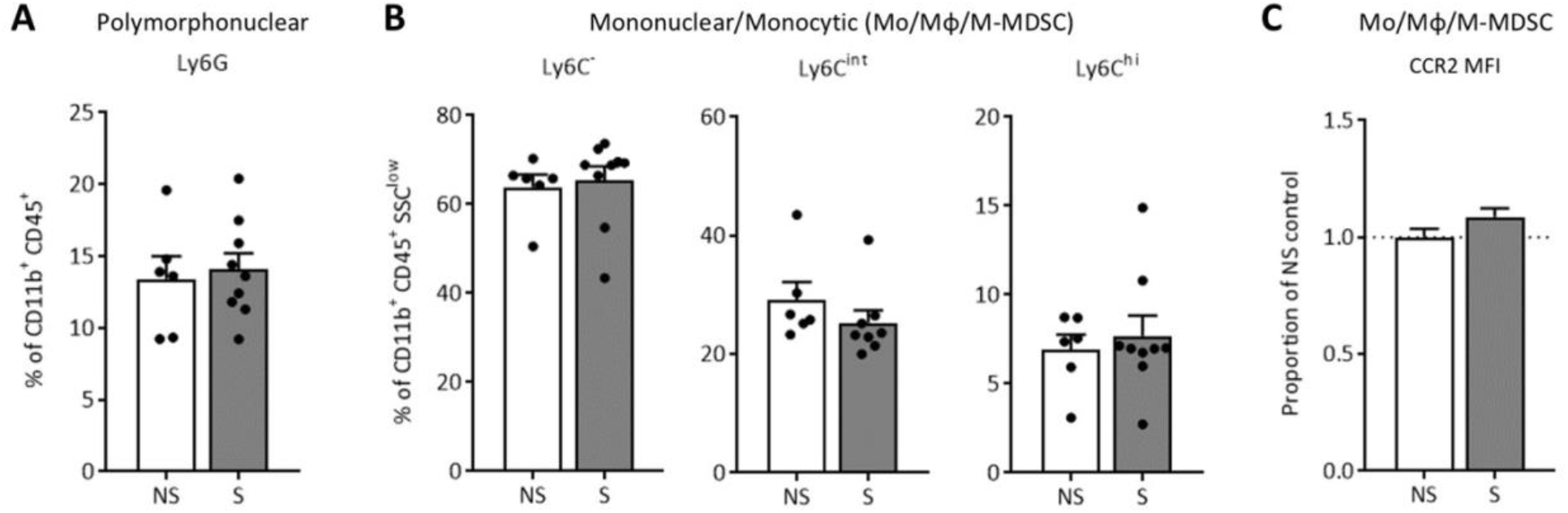
GD17 uterine CD11b^+^CD45^+^ leukocytes appear to be stress resistant. Flow cytometric analysis of CD11b^+^CD45^+^ cells revealed that stress did not impact uterine **(A)** polymorphonuclear cell percentages (p = 0.74; neutrophils or PMN-MDSCs), or **(B)** percent of SSC^low^ mononuclear cells within the three phenotypes (Ly6C^-^ alternative M2, p = 0.77; Ly6C^int^ transitional, p = 0.30; or Ly6C^hi^ classical M1, p = 0.62). **(C)** CCR2 MFI within uterine Mo/Mϕ/M-MDSCs isolated from restrained dams did not differ from controls (p = 0.13). Data are mean ± SEM, with each dot representing a single dam. NS = non-stressed, n = 6; S = stressed, n = 9; PMN-MDSCs = polymorphonuclear myeloid-derived suppressor cells; Mo/Mϕ/M-MDSCs = monocyte/macrophage/monocytic myeloid-derived suppressor cells; MFI = median fluorescence intensity.

### 2.4. Prenatal stress decreases CD11b+ cell density in the GD17 placenta, but polymorphonuclear and mononuclear leukocyte proportions are protected

CD11b^+^CD45^+^ leukocytes within placental tissue were examined at GD11 (after two two-hour sessions of maternal restraint stress) and at GD17 (24 h after the conclusion of the seven-day restraint stress paradigm) using flow cytometry. At GD11, the percent of CD11b^+^CD45^+^ cells classified as Ly6G^+^ (polymorphonuclear) or SSC^low^ on the Ly6C spectrum (mononuclear; Mo/Mϕ/M-MDSCs) did not differ with prenatal stress and displayed high variability (**Supp. Fig. 1**). At GD17, cell counts (cells/mg of tissue) of CD11b^+^CD45^+^Ly6G^+^ polymorphonuclear cells (neutrophils or PMN-MDSCs) did not differ (**Fig. 4A**), but a substantial decrease in overall CD11b^+^CD45^+^SSC^low^ mononuclear cell counts (Mo/Mϕ/M-MDSCs) was evident and was reflected across all Ly6C cell phenotypes (**Fig. 4B**). The percent of CD11b^+^CD45^+^ cells identified as polymorphonuclear Ly6G^+^ (neutrophils or PMN-MDSCs; **Fig. 4C**), or mononuclear Ly6C^-^ alternative M2, Ly6C^int^ transitional, and Ly6C^hi^ classical M1 (Mo/Mϕ/M-MDSCs; **Fig. 4D**) did not differ, indicating that cell proportions remain relatively stable despite an overall reduction in total mononuclear cell numbers. Like at GD11, CD11b^+^CD45^+^SSC^low^ Mo/Mϕ/M-MDSCs within GD17 placental tissue did not differ in expression of CCR2 (**Fig. 4E**). As gestation progressed from GD11 to GD17, leukocyte proportions shifted independent of prenatal stress, as evidenced by an increase in Ly6G^+^ and Ly6C^hi^ cells, and a resultant reduction in transitional Ly6C^int^ cells, at GD17 compared to GD11 (**Supp. Fig. 2**).

**Fig. 4.**
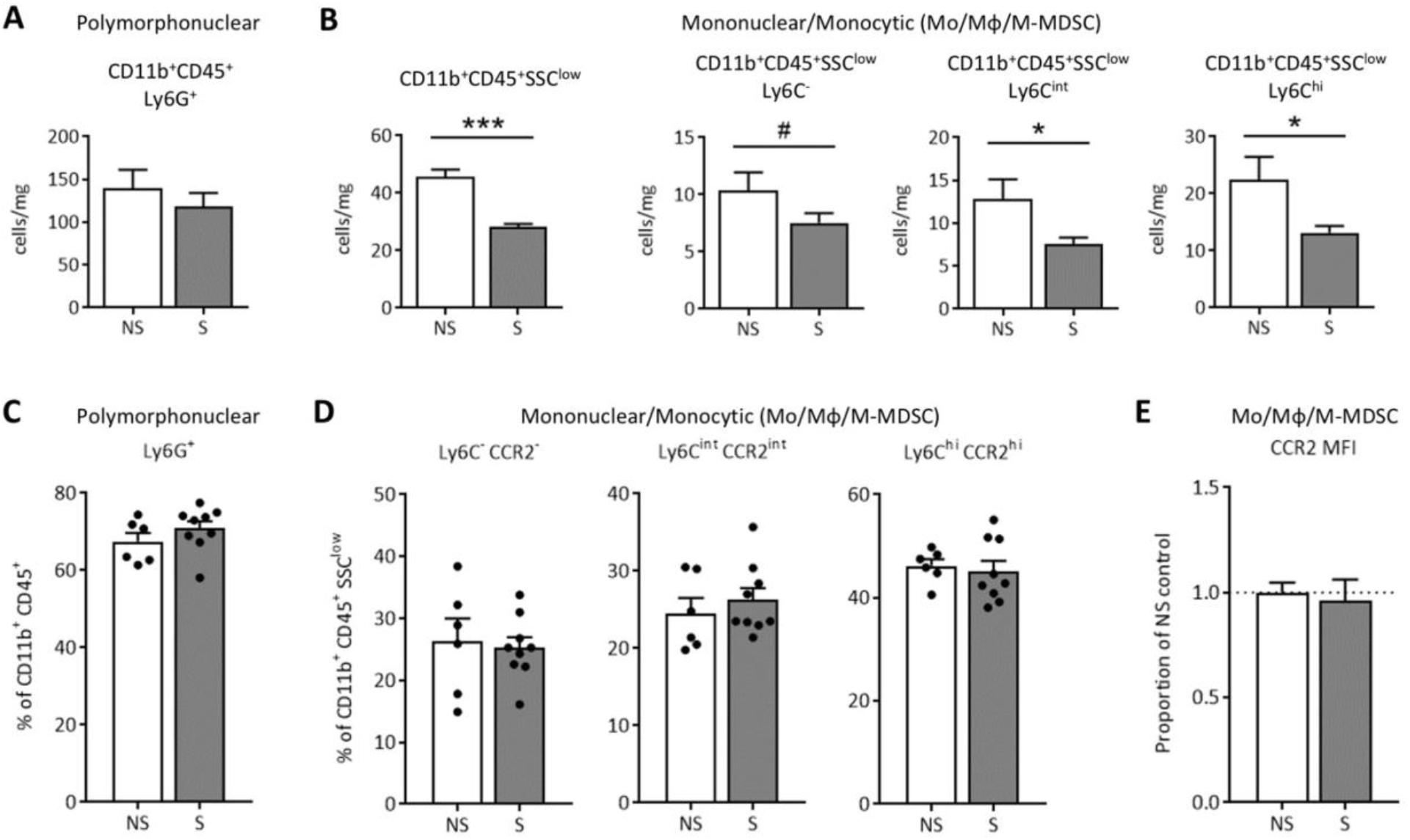
Placental CD11b^+^CD45^+^SSC^low^ leukocyte density decreases with prenatal stress at GD17, but cell proportions remain stable. Flow cytometric analysis of CD11b^+^CD45^+^ cells revealed **(A)** no change in polymorphonuclear Ly6G^+^ cell numbers (p = 0.45) but **(B)** a dramatic decrease in total SSC^low^ mononuclear cells (p < 0.0001) due to prenatal stress that was apparent across all three Ly6C subtypes (Ly6C^-^ alternative M2, p = 0.10; Ly6C^int^ transitional, p = 0.04; and Ly6C^hi^ classical M1, p = 0.04). However, percent of placental CD11b^+^CD45^+^ cells classified as **(C)** Ly6G^+^ polymorphonuclear cells (neutrophils or PMN-MDSCs; p = 0.28) and **(D)** SSC^low^ mononuclear cells did not significantly differ across treatment (Ly6C^-^ alternative M2, p = 0.77; Ly6C^int^ transitional, p = 0.50; and Ly6C^hi^ classical M1, p = 0.71). Likewise, **(E)** CCR2 median fluorescence intensity did not shift among the Mo/Mϕ/M-MDSC population (p = 0.73). Data are mean ± SEM, with each dot representing a pooled sample (containing a minimum of 2 placentas) from 1 litter. NS = non-stressed, n = 6, S = stressed, n = 9; Mo/Mϕ/M-MDSCs = monocytes/ macrophages/monocytic myeloid-derived suppressor cells; MFI = median fluorescence intensity. *** = p <0.001, * = p < 0.05, # = p ≤ 0.10.

### 2.5. Psychological stress reduces the abundance of multiple bacterial groups of the gastrointestinal microbiome

The microbiome has long been implicated in modulating host immunity and stress responses. Thus, potential stressor-induced effects on the maternal gastrointestinal microbial community structure and function were examined. Colon contents from GD17 stressed and non-stressed dams were analyzed through DNA extraction, shotgun metagenomic sequencing, and both genome-resolved and read-based metagenomic analyses. Using LefSE, we identified multiple metagenome-assembled genomes (MAGs) that significantly associated with either stress or non-stress (p < 0.05, LDA > 2.0). MAGs increased in non-stressed dams were members of the *Atopobiaceae* family, the *Acetatifactor* genus, the *UBA7173* genus of the *Muribaculaceae* family, multiple members of the *Lachospiraceae* family including *Eubacterium* (formerly of the *Clostridiales* order, now classified under *Lachnospirales^44^)* and *Parasutterella excrementihominis* (**Fig. 5A**). MAGs that were increased in stressed dams were a member of the *UBA9502* genus and the *Lachnospiraceae* family and a member of the *Oscillibacter* genus (Fig. 5A). Further examination of *Parasutterella excrementihominis* genome coverage revealed significant differences between groups as per Kruskal Wallis and Wilcoxon Signed Rank tests (**Fig. 5B**, p = 0.006, LDA = 2.94).

**Fig. 5.**
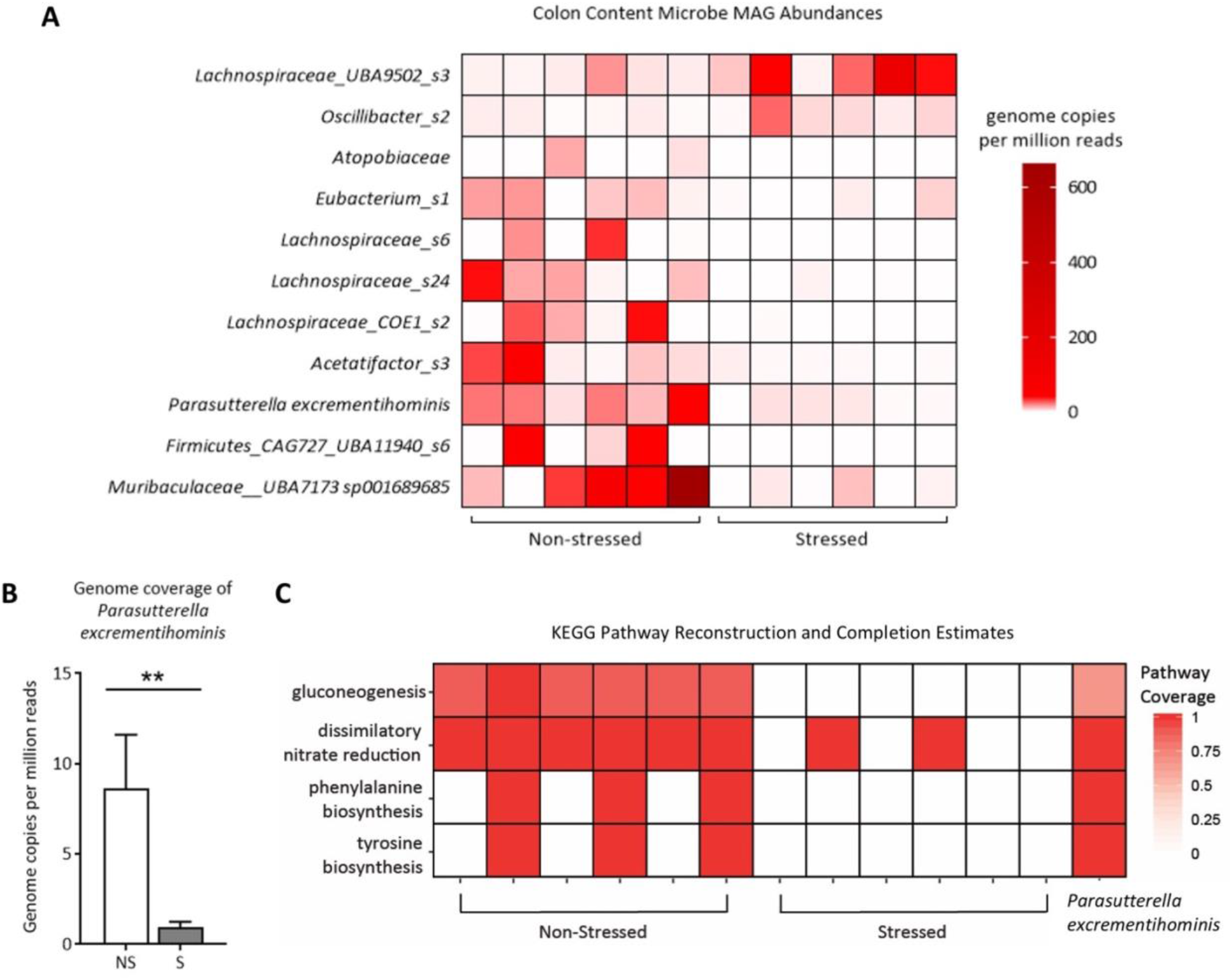
GD17 colonic microbiota metagenome analyses reveal restricted bacterial abundances and metabolic pathways due to psychological stress. **(A)** Differentially abundant metagenome-assembled genomes (MAGs) in non-stressed and stressed samples, as identified by LefSe, using genome copies per million reads. Taxonomy is GTDB taxonomy with the lowest possible classification listed. Some MAGs have higher level taxonomy listed for easier identification. **(B)** *Parasutterella excrementihominis* genome copies per million reads (mean ± SEM; Kruskal Wallis and Wilcoxon Signed Rank tests, p = 0.006). **(C)** KEGG pathway completion for select KEGG pathways in whole sample assemblies that displayed presence/absence differences between sample groups using KEGGDecoder. Samples are grouped by non-stress and stress, with the KEGG pathway completions for the consensus *Parasutterella excrementihominis* MAG on the far right. NS = non-stressed, S = stressed; n = 6/group; ** = p < 0.01.

### 2.6. Psychological stress alters microbial gene abundances and pathways associated with *Parasutterella excrementihominis*

Given that immunity and inflammatory suppression were prime targets of stressor activation, microbial genes that may play a role in host immune function were examined. Multiple gene functional groups were elevated in non-stressed dams compared to stressed dams including: *ttrS* (sensor kinase for tetrathionate two-component regulatory system, K13040, p < 0.0001), *dcuCD* (C4-dicarboxylate transporter of the DcuC family, K00326, p < 0.0001), and *frdA* (fumarate reductase flavoprotein subunit A, K00244, p < 0.0001). Fumarate reductase is implicated in the production of succinate, which is typically linked to inflammation in the gut during stress but may be involved in enabling and sustaining the growth of commensal fermentative gut microbes^45–48^. As such, we further identified MAGs capable of succinate production in both stressed and non-stressed dams. Three (of five) *Parasutterella excrementihominis* MAGs in non-stressed dams contained genes encoding for fumarate reductase (K00244/K00245/K00246, missing K00247), notably the only MAGs with these particular genes (Supporting File A). However, an alternative enzyme encoding for the conversion of fumarate to succinate, succinate dehydrogenase/fumarate reductase (K00239/K00240/K00241 missing K00242) were present in many other MAGs. This enzyme was present in members of the *Dehalobacteriia* class, the *Muribaculaceae* family, *Akkermansia municiphila, Bacteriodes thetaiotamicron*, and *Alistipes* species, distributed throughout both non-stressed and stressed dams. Within MAGs capable of converting fumarate to succinate, *Parasutterella excrementihominis, Akkermansia municiphila, Bacteroides thetaiotamicron*, and *Dehalobacteriia* contained genes associated with conversion of succinate to succinyl-CoA (K01902/K1903 or K18118). While succinate production appears widely distributed amongst taxa and in both sample groups, the fumarate reductase gene subunit A (frdA, K00244) was significantly higher in non-stressed dams compared to stressed dams, and since *Parasutterella excrementihominis* was the sole taxa to contain this gene, we surmise it is the microbe responsible for the difference.

We further investigated putative metabolic differences in dam colonic content based on KEGG pathways reconstructed from microbial genomes as a function of stress or non-stress. Pathways for gluconeogenesis, dissimilatory nitrate reduction, and phenylalanine and tyrosine biosynthesis were absent in the majority of stressed dams, and their presence was numerically increased in non-stressed dams (**Fig. 5C**). Interestingly, in non-stressed dams, all of these pathways were identified in *Parasutterella excrementihominis*, but were absent in other differentially abundant MAGs, indicating that the absence of functional genes corresponding to these pathways is likely the result of decreased *Parasutterella excrementihominis* in stressed dams. We further confirmed the absence of key genes required for gluconeogenesis and tyrosine biosynthesis in all other MAGs. Genes necessary for dissimilatory nitrate reduction were present in one MAG in one stressed sample (of the *Adlercruetzia* genus), but otherwise, *Parasutterella excrementihominis* contained the only dissimilatory nitrate reduction genes in these samples.

### 2.7. Maternal intestinal gene expression is only moderately influenced by psychological stress

Given that restraint stress altered the colonic microbiome, we hypothesized that stressor-induced differential expression may be present in intestinal genes involved in inflammation and barrier integrity. Gene expression analysis of tight junction proteins and alpha-defensins revealed only a decrease in claudin 5 (*CLDN5*) and alpha-defensin 2 (*DEFA2*) in colon tissue (expression of genes associated with barrier integrity and immune response are listed in **Supp. Table 5**). Changes in colonic toll-like receptor 2 and 4 (*TLR2* and *TLR4*) gene expression that did not reach significance (**Supp. Table 5**). While a disruption in colonic tight junctions may indicate barrier disruption, serum LPS-binding protein was not impacted by stress (data not shown; p = 0.82; NS: 52.1 ± 5.3 ng/mL, n = 14, S: 53.7 ± 4.5 ng/mL, n = 13). Although expression of *CLDN5* numerically decreased within the ileum (p = 0.13), no gene expression changes reached significance (**Supp. Table 5**).

## 3. Discussion

Exposure to prenatal stress can have life-long negative impacts on offspring mental health and wellbeing, and the mechanisms underlying these risks have yet to be completely defined. Maternal immune and glucocorticoid responses have repeatedly been demonstrated to be two of the primary drivers of poor infant outcomes^49,50^, although variability in stress severity and duration of exposure make it difficult to pinpoint critical activation thresholds of these cellular and molecular signaling pathways. Expanding investigations of the maternal microbiome during stress and gestation also highlight microbes as primary mediators of immune status and infant neurodevelopment^51^. Our murine model of prenatal stress defines critical maternal glucocorticoid, immune, and microbial alterations that may underlie the negative impacts of psychological stress during gestation. In particular, our data indicate that nuanced and complex interactions between glucocorticoid and microbial signals drive a unique maternal immune profile that is in contrast with documented fetal neuroinflammation.

Gestational stressor paradigms are often selected for *not* influencing maternal weight gain and litter size^52^, so as to limit confounding factors. In our model, restraint stress did not impact overall litter size or number of resorptions. Pregnant dam body weight and litter size at GD17 also did not differ between treatment groups, which is consistent with our previous reports^25^ and in agreement with other rodent models of prenatal stress^52–54^. However, when weight gain was normalized to litter size, restraint stress was revealed to restrict proper increases in maternal weight over the course of pregnancy. To our knowledge, this is the first time such a comparison has been conducted within the prenatal stress animal literature and may reveal a “fetal sparing” effect.

An increase in circulating pro-inflammatory cytokines is one of the hallmarks of pregnancy complications^55,56^ and can be detected in pregnant women experiencing psychological stress^50,57,58^. In animal models, systemic administration of particular cytokines in circulation, such as IL-6 and IL-1β, is sufficient to recapitulate specific offspring neuroimmune abnormalities seen with prenatal stress^59,60^. Interestingly, while psychological stress is thought to induce cytokine production in animal models (and is frequently described in placental and offspring brain tissue, including by our group^25,42^), few, if any, studies have reported maternal circulating cytokine levels. Here, we found a decrease in IL-5 and a trending decrease in CCL2 in maternal circulation, and none of the suspected increases in IL-6 and IL-1β. If anything, a *reduction* in maternal splenic production of IL-1β would indicate the opposite in our model, which is consistent with the immunosuppressive phenotype induced by restraint stress^61,62^. An activation of the HPA axis and increased circulating corticosterone observed here, which is continually described in both humans and animals during prenatal stress^49,63^, contributes to this immunosuppressive phenotype^62^.

Animal modeling of prenatal stress has revealed that maternal inflammatory pathways may be responsible for offspring behavioral abnormalities^59,60,64^, yet potential contribution of disrupted leukocyte populations in maternal circulation or at the maternal-fetal interface have received little attention in this context. Some have demonstrated conflicting results of prenatal social stress on blood monocyte counts in rats^54^ and swine^65^, and no change in neutrophils. At the maternal-fetal interface, acute restraint stress on GD1 has been shown to disrupt endometrial lymphocyte concentrations and attenuate proliferation and cytokine secretion^66^. Shifts in intrauterine leukocyte populations, driven by localized cytokine signaling^67,68^, have been demonstrated in animal models of direct intrauterine inflammation^69^. However, the decrease in or absence of leukocyte recruitment in our study (in the placenta and uterus, respectively) indicates that prenatal stress is unique and distinct from these models. To our knowledge, this is the first time that neutrophils and monocytes have been assessed in circulation and at reproductive sites during prenatal restraint stress. Furthermore, unlike other models^68^, mRNA transcripts of inflammatory signaling molecules within uterine tissue were not altered in our model. Even still, our previous observations demonstrate significant impacts of gestational restraint stress on offspring inflammation and behavior^25,42^, which are largely CCL2-dependent^43^. Altered production and expression of CCL2 and CCR2 in both the fetal brain and placenta coincide with pro-inflammatory signatures within these tissues and appear to drive offspring sociability^43^. Therefore, it is possible that more nuanced inflammatory signaling patterns (involving CCL2-CCR2 chemokine signaling) are occurring systemically and at the maternal-fetal interface that are not reflected in CD11b^+^CD45^+^ cell proportions here. Future investigations will focus on untangling systemic and localized CCL2-dependent immune mechanisms across the duration of restraint stress (GD10-GD16), which might lead to altered fetal neurodevelopment in our model.

As systemic indicators of the maternal immune response to gestational stress are incompletely defined within animal models, our data reinforce the need for comprehensive exploration of these pathways across common prenatal stress models. Indeed, the unique immune profile of pregnancy alone may produce a conflicting phenotype when compared to non-pregnant animals exposed to repeated restraint stress, which exhibit reductions in circulating leukocytes^61,70^ and elevations in IL-6^71^. Type and duration (chronic vs. acute) of a stressor paradigm also has significant impacts on glucocorticoid and immune variables^61^, and gestational timing must be considered^72^. Thus, indications of lasting or acute maternal immunosuppression during prenatal stress, as we report, need to be reconciled with inflammatory signaling within the fetal brain and placenta that appears to be driving offspring outcomes.

While our group^25^ and others^26^ have shown that prenatal stress can alter maternal fecal microbial populations, the intestinal microbial community has not yet been fully explored in the context of prenatal stress. Largely, our data stand in contrast to recent studies that have highlighted stark decreases in immune-modulatory bacterial groups such as *Lactobacillus*^73,74^, as well as exacerbations in inflammatory severity upon exposure to stress^75,76^. The previously accepted belief was that psychological stress exacerbated GI inflammation, an outcome that can also be observed clinically given the propensity for stressful life events to be associated with relapse in inflammatory bowel disease^77^. However, our results indicate that in pregnant dams, stressor exposure can also act in an immunosuppressive context within the microbial metagenome. Stressor-induced reductions in *ttrS*, the sensor kinase for the tetrathionate two-component regulatory system, and *dcuCD* and *frdA*, both of which have involvement in succinate production, indicate that suppression of microbial genes may contribute to reduced inflammatory responses in the host. Tetrathionate is closely associated with intestinal epithelial inflammation and gives pathogenic *Salmonella* an added fitness niche for growth during GI colonization^78^, while succinate can induce IL-1β and activate dendritic cells in a pro-inflammatory manner^47,48^.

That these genes were reduced points to a concomitant reduction in immune-activating microbes. Indeed, multiple microbial groups were reduced by stress, including members of the *Clostridiales* order, members of the *Lachnospirales* order including a strain of *Eubacterium*, and the species *Parasutterella excrementihominis*. A previously studied *Eubacterium* strain (*E. lentum)* was capable of dehydroxylating cortisol to 21-deoxycortisol^79^, indicating the potential that *Eubacterium* here may participate in cortisol metabolism. *Parasutterella excrementihominis* is of particular interest, as it is a known health-associated bacterial group^45^. Though further examination of the integral nature of *Parasutterella* in both dam as well as offspring health is outside of the purview of this particular study, our data suggest that *Parasutterella excrementihominis* plays a key role in maintaining host immunity during late pregnancy. Indeed, Ju et al. (2019) have identified a central role for *Parasutterella* in gut health, demonstrating that this genus is not only a known colonizer of the mouse gut, but also a producer of succinate. Succinate has been previously shown to indirectly support the growth of fermenters such those in *Clostridiales*, preventing infection from pathogens in neonatal mice^45,46^. Future studies will investigate how succinate may be involved in immune and inflammatory regulation during pregnancy.

Studies show that lower levels of *Parasutterella* are associated with high-anxiety in pregnant human mothers^80^ and in stressor-exposed male mice^81^, indicating that there is universality in stressor sensitivity for this particular microbial group. The reason for these targeted reductions is likely multi-layered. Psychological stress increases gut motility^82,83^, which may increase shear stress on mucosal-adhered microbial groups. In addition, given the fact that the microbiota exist in climax communities, alterations in mucosal immune function, nutrient availability, or epithelial barrier stability could all have drastic effects on microbiome structure^84,85^. Our study and others depict an environment wherein stress can be both pro- and anti-inflammatory^76,86^, systemically and at the gastrointestinal epithelium. Resultant outcomes include relaxed or elevated macrophage trafficking and antigenic presentation activity^87–90^ and barrier dysfunction^91,92^, all of which may act profoundly upon microbial abundances. However, an understanding of why *Parasutterella* is a frequent target is presently incomplete. Potentially, *Parasutterella’s* unique inability to metabolize carbohydrates and utilization of amino acids as a primary energy source may leave it more susceptible to stressor-induced fluctuations in substrate availability^45^, though this must still be tested.

*Parasutterella* is a key player in multiple GI metabolic pathways, including tryptophan, tyrosine and bile acid metabolism^45^. Here, multiple metabolic pathways were almost completely absent in stressor-exposed pregnant dams, including gluconeogenesis, dissimilatory nitrate reduction, and both the phenylalanine and tyrosine biosynthesis pathways. All of these pathways are present in *Parasutterella excrementihominis*, and we confirmed absence of gluconeogenesis, dissimilatory nitrate reduction, and tyrosine biosynthesis in other MAGs from non-stressed dams. We further surmise that it is because of the stressor-induced reduction of *Parasutterella* that we are unable to observe these particular pathways in stressor-exposed dams. These findings further strengthen the argument that *Parasutterella* is integral to proper gut health during pregnancy. Our stressor paradigm continues into late gestation, which is characterized by a pro-inflammatory shift in order to support labor and delivery^4^, which may be reflected in the increased classical Ly6C^hi^ placental mononuclear cells at GD17 compared to GD11. Thus, a reduction in microbial inflammatory genes, including those associated with succinate, may be detrimental both for appropriate microbial transfer from mother to infant^29,93^ and for necessary infant immune education^94^. Long-term effects of maternal stress upon offspring notably include microbiome shifts^25,26,42^, but also spread into basic immune function, including elevated inflammatory output and a dysregulated adaptive immune response^95,96^. There is the strong potential that *Parasutterella* may mediate proper metabolic function in late pregnancy and even early immune development in the offspring; future studies by our group will delve deeper into how *Parasutterella* and its production of metabolites (i.e. succinate) may fortify either maternal or infant health.

The unique maternal immune and microbial profiles unveiled in our study indicate that repeated restraint stress imparts complex immunomodulatory effects during the mid-to-late gestational period. Stressor-induced glucocorticoid signals appear to induce immune aberrations systemically and at the placenta. A concomitant disruption in GI microbial communities not only mirrors a unique immunosuppressive functional phenotype but may also feedback upon and contribute to localized and systemic immune capacity. Furthermore, our data reveal a unique maternal immune phenotype during prenatal stress that does not mirror the inflammatory responses elicited in the fetal brain and placenta, indicating that the intrauterine immune signaling patterns driving fetal outcomes are more intricate and nuanced in this context. These exciting findings expand our understanding of the potential impacts of prenatal psychological stress on the developing fetus, and also provide several avenues for developing non-invasive therapeutic strategies that could be applied prenatally.

## 4. Materials and Methods

### 4.1. Animals

Nulliparous adult C57BL/6J female mice were acquired from Jackson Laboratories (Bar Harbor, ME) at 10-weeks of age and acclimated to the vivarium at The Ohio State University Wexner Medical Center for a minimum of one week. Females were bred over two nights and pregnancy was verified by the presence of a vaginal plug (designated as GD1). Pregnant females were randomly assigned to the experimental stress group or the non-stress control group, with the stress group undergoing repeated restraint stress from GD10 to GD16, as previously described^25,42^. Briefly, pregnant dams were placed in a perforated 50 mL conical for two hours each day between the hours of 09:00 to 12:00 for seven consecutive days; non-stressed control dams were left undisturbed. Pregnant dams were sacrificed at GD17, 24 h after the conclusion of the stressor paradigm, and maternal and placental tissues were collected (the study design is depicted in **Fig. 1A**). A subset of animals was sacrificed at GD11, immediately following the second implementation of restraint stress, to examine placental leukocytes. For all examinations of placental leukocytes, an average of two placentas per litter were included (balanced for sex), with no more than four placentas used per litter. At sacrifice, litter size and fetal resorptions were determined. All procedures were in accordance with and approved under the Institutional Animal Care and Use Committee at The Ohio State University.

### 4.2. Tissue collection

Pregnant dams were euthanized by inhalation of CO_2_. Whole blood was collected through cardiac puncture into EDTA-coated tubes (for plasma collection and whole blood flow cytometry; kept on ice) or into non-coated tubes (for serum collection; kept at room temperature). The uterus was sterilely excised through cesarean section and individual fetuses and placentas were dissected out. Maternal brains were excised and micro-dissected using a mouse brain matrix to identify and collect specific regions, as previously described^25,42^. Tissues were either snap frozen and stored at −80°C until further processing (for gene and protein analyses) or placed into sterile ice-cold HBSS (without Ca^2+^ or Mg^2+^) and stored on ice until further processing (for leukocyte isolation and flow cytometric analyses). Serum tubes were allowed to clot at room temperature for a minimum of 30 min; serum and plasma blood tubes were spun at 13,300 rpm at 4°C for 15 min and then aliquoted and stored at −80°C until further processing.

### 4.3. Protein Isolation

Protein was isolated from half of each spleen and uteri, which were weighed prior to protein extraction (the other halves were used for gene expression analyses). Using sonication at 40% amp, each sample was homogenized in 2 mL of lysis buffer, containing 100:1 T-PER™ Tissue Protein Extraction Reagent (Thermo Scientific™, Waltham, MA) and Halt™ Protease and Phosphatase Inhibitor Single-Use Cocktail (100X; Thermo Scientific™). Homogenized samples were incubated at 4°C for 30 min and then centrifuged at 13,300 rpm at 4°C for 15 min. Supernatant was aliquoted and immediately assayed using the Pierce™ BCA Protein Assay kit (Thermo Scientific™), according to manufacturer’s protocol, to determine protein concentration for each sample. Protein concentration was then normalized to 1.8 mg/mL for each sample before utilization in ELISAs (below).

### 4.4. ELISAs

For measuring corticosterone, serum was diluted 1:10 in diluent and run in duplicate using Cayman Chemical (Ann Arbor, MI) Corticosterone ELISA kit, following manufacturer’s protocol. Protein lysates were assayed in duplicate for cytokines IL-1β, TNFα, and CCL2 using Mouse IL-1 beta/IL-1F2, TNF-alpha, and CCL2/JE/MCP-1 DuoSet ELISA kits (R&D Systems™, Minneapolis, MN) according to manufacturer’s protocol; data were normalized to tissue weight (mg) and are presented as proportion of non-stressed control. Plasma was assayed in duplicate for CCL2 using the same CCL2/JE/MCP-1 DuoSet ELISA kit (R&D Systems™) according to manufacturer’s protocol. Using the MSD^®^ Multi-Spot Assay System Proinflammatory Panel 1 (mouse) V-Plex kit, serum was diluted 1:1 with Diluent 41 and assayed according to manufacturer’s instructions using Alternate Protocol 1, Extended Incubation (Meso Scale Discovery^®^, Rockville, MD). For measuring LPS-binding protein, serum was diluted 1:5 in diluent buffer and run in duplicate using Mouse LBP ELISA Kit (Abcam, Cambridge, MA) according to manufacturer’s instructions.

### 4.5. Quantitative real-time PCR

RNA was isolated using 1 mL TRIzol™ Reagent (Invitrogen™, Carlsbad, CA) per sample following the TRI Reagent^®^ Protocol. Applied Biosystems™ (Foster City, CA) High-Capacity cDNA Reverse Transcription Kit was used to synthesize cDNA as per manufacturer’s instructions. PCR was performed in duplicate using TaqMan™ Fast Advanced Master Mix (Applied Biosystems™) in 384-well plates, as per instructions, and assayed on a QuantStudio™ 3 Real-Time PCR System machine (Applied Biosystems™). Gene primer information is presented in **Supp. Table 6**. Data were analyzed using the 2^-ΔΔCt^ method with housekeeping gene *RPL19* and are presented as relative expression compared to the non-stressed control group.

### 4.6. Leukocyte characterization

The following rat anti-mouse antibodies were used for leukocyte analysis within each tissue: purified CD16/CD32 Fc blocking antibody (clone 2.4G2; BD Pharmingen™, San Jose, CA), PerCP-Cy™5.5 Ly6G antibody (clone 1A8; BD Pharmingen™), V450 CD45 (clone 30-F11; BD Horizon™, San Jose, CA), APC CD11b (clone M1/70; eBioscience™, San Diego, CA), PE-Cyanine7 Ly6C (clone HK1.4; eBioscience™), PE CCR2 (clone 475301, R&D Systems™). UltraComp eBeads™ Compensation Beads (Invitrogen™) were used for single-stain compensation controls as per manufacturer’s instructions.

#### 4.6.1. Whole blood

50 μL of whole blood collected into EDTA-coated tubes was used for circulating leukocyte analysis. Samples were incubated with fluorescent antibodies (listed above) for 30 min on ice protected from light, then incubated with 1 mL RBC Lysis Buffer for 15 min. Lysis buffer was quenched with 3 mL sterile PBS and samples were centrifuged at 1200 rpm at 4 °C for 6 min and supernatant was removed. Cells were fixed in 10% neutral buffered formalin (NBF) for 10 min and then resuspended in FACS buffer (1% BSA and 2mM EDTA in sterile PBS). 100 μL of 123count eBeads™ (Invitrogen™) were added to each sample prior to analysis.

#### 4.6.2. Uterus and placenta

Uterine and placental tissues were transferred from ice-cold HBSS into 5 mL enzyme digestion media (10% FBS, 10mM HEPES, 6 U DNase I, and 2mg/mL Collagenase A in sterile HBSS with Ca^2+^ and Mg^2+^) and minced before incubating with gentle shaking at 37°C for 30 min. Samples were then passed through a 40 μm cell strainer to create a single-cell suspension and rinsed with FACS buffer before pelleting at 1,250 x g at 4°C for 10 min and then resuspending in 90 μL FACS buffer. Cells were then incubated with CD11b MicroBeads (mouse/human; Miltenyi Biotec Inc., Auburn, CA) and passed through MACS^®^ LS magnetic columns for positive selection of CD11b^+^ cells, as per manufacturer’s instructions. Cells were then incubated with fluorescent antibodies (listed above) for 30 min on ice protected from light, and then fixed in 10% NBF for 10 min. Stained fixed cells were resuspended in FACS buffer and 100 μL of 123count eBeads™ (Invitrogen™) were added to placental samples prior to analysis.

#### 4.6.3. Flow cytometry

Stained fixed cells were all analyzed on a BD FACSCalibur™ cytometer (BD Biosciences™, San Jose, CA) using FlowJo™ 10 software. Cells were gated on forward-(FSC) and side-scatter (SSC) properties to identify live leukocytes, and then gated as CD45^+^ and CD11b^+^. Polymorphonuclear cells were identified as CD11b^+^CD45^+^ cells with a mid-to-high SSC, and then gated on Ly6G. Mononuclear cells were identified as CD11b^+^CD45^+^ with a low SSC, and then gated on Ly6C and CCR2. A massed unstained control sample (for each respective tissue) was used to gate negative and positive CCR2 expression within CD11b^+^CD45^+^SSC^low^ mononuclear cells, where any event beyond the PE autofluorescence boundary was considered CCR2^+^. Median fluorescence intensity (MFI) of CCR2 is presented as proportion of NS control. Cell counts were calculated based on instructions in the 123count eBeads™ protocol (Invitrogen™) and then normalized per μL of whole blood or mg of tissue.

### 4.7. DNA sequencing, quality control, and metagenomic analysis of colon contents

Twelve colon content samples at GD17 were selected for DNA extraction and sequencing, six from stressed dams and six from non-stressed dams. DNA was extracted using a QIAGEN PowerFecal extraction kit (QIAGEN Inc., Germantown, MD), according to the manufacturer’s instructions. DNA was quantified and quality checked using a Qubit 4 Fluorometer and a Nanodrop 2000 Spectrophotometer (Thermo Scientific™). DNA metagenomic libraries were generated using the KAPA Library System (per manufacturer’s instructions). After quality control measures, libraries were sequenced using the Illumina NovaSeq SP flow cell (paired-end 150 bp) at the OSUCCC Genomics Shared Resource. All paired end raw reads were trimmed using TrimGalore (http://www.bioinformatics.babraham.ac.uk/projects/trim_galore/) and bmtagger was used to remove host reads by mapping against the mm10 mouse genome. Samples were assembled individually with MEGAHIT^97^ and assembly quality and read alignment rates were assessed using metaQUAST^98^ and Bowtie2^99^. Scaffolds ≥ 1 kb from the final assembly were binned using MetaBAT2^100^ and CONCOCT^101^ followed by bin consolidation, refinement, and reassembly using metaWRAP^102^ to identify metagenome-assembled-genome bins (MAGs). We retained MAGs that were 75% complete with less than 5% contamination, according to CheckM^103^. Taxonomy was inferred using GTDB-Tk^44^. Consensus MAGs were obtained using dRep^104^ to dereplicate MAGs across all 12 samples, with default settings applied. Bin quantification per sample (genome copies per million reads) was performed on dereplicated consensus MAGs in metaWRAP using Salmon^105^. Using a genome-resolved metagenomic approach with individual sample assembly and binning, 591 medium to high quality (> 75% completion with < 5% contamination per CheckM^103^) metagenome-assembled genomes (MAGs) were recovered. After dereplication of these MAGs, 232 consensus MAGs were retained and used for downstream analyses. The coverage of these MAGs was calculated using the bin quantification approach stated above. Differentially abundant MAGs between sample groups were identified using the results of the bin quantification (genome copies per million reads) in LefSe, with default settings (p <0.05, LDA > 2.0)^106^. Genes within assemblies and in MAGs were identified by translating nucleotide sequences into protein sequences using Prodigal^107^, then proteins were annotated using hidden markov models via KOFamScan^108^. Pathway coverage (0-1) was estimated using KEGGDecoder^109^ with the output from KOFamScan. We used both DESeq2 (in R v. 3.6.3)^110^ and LefSe to identify differentially abundant genes between sample groups, with default settings and significance of p < 0.05.

To confirm the findings from the genome-resolved metagenomic approach listed above, we also performed a read-based metagenomics approach and analyzed the trimmed reads (paired ends were merged) using HUMAnN 2.0^111^ and MetaPhlAn2^112^ using the default settings. Gene families, bacterial taxa, and pathway abundances that were significantly different between sample groups were assessed using LefSe and MAASLIN (http://huttenhower.sph.harvard.edu/maaslin2).

The raw reads from this study are publicly available under the BioProject PRJNA632941, with sample accessions available in Supporting File A.

### 4.8. Statistics

All data (except metagenomic analyses, as stated above) were analyzed using GraphPad Prism Software (San Diego, CA). Non-stressed and stressed groups were compared using unpaired parametric t tests, with Welch’s correction when variances were unequal. Two-way ANOVA was used when comparing multiple variables (i.e. prenatal treatment x gestational day when comparing placental leukocytes at GD11 and GD17). Linear regression analysis was used to plot gestational weight gain against litter size and to test whether slopes and intercepts were significantly different between prenatal treatment groups. For all tests, significance was set at α = 0.05, and outliers were identified and removed using the ROUT method at Q = 1%.

## 6. Acknowledgements

This work was supported by K08 MH112892, R21 MH117552, and start-up funds from the Ohio State University to T.L.G., T32 DE1432017 to A.M.A., and T32 NS105864 to H.J.C. We would like to acknowledge Sydney Schiff and Amber Dalal for their contributions.

## Supplemental Materials

### Supporting File A

Uploaded as Excel spreadsheet. Metagenomic accession numbers and assembly statistics, MetaPhlAn2 taxonomy, KOFAMScan gene counts, Consensus bin genome copies per million reads in each sample, succinate metabolic gene presence/absence in all MAGs, and GTDBTk taxonomy for both full and consensus MAG sets.

**Supp. Fig. 1.**
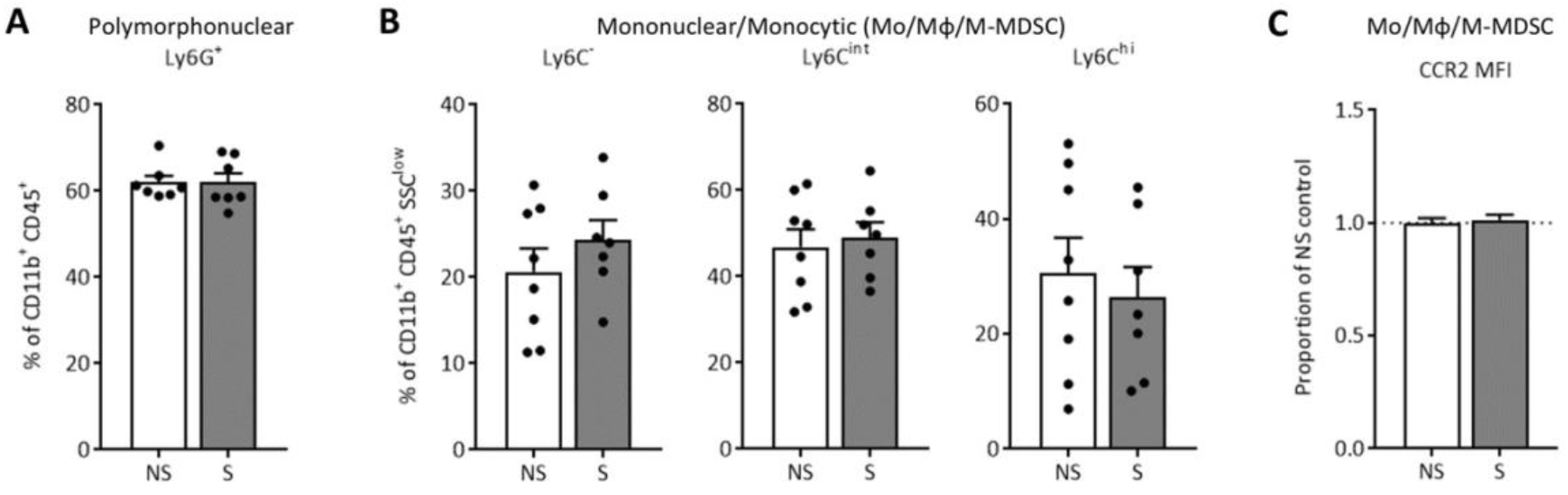
GD11 placental leukocyte populations are protected from prenatal stress. Investigation of GD11 placental tissue indicated that CDllb*CD45* polymorphonuclear (neutrophils or PMN-MDSCs) and SSC^low^ mononuclear (Mo/Mϕ/M-MDSCs) cell populations are highly variable at GD11, and that two days of maternal restraint stress does not shift CCR2 expression in Mo/Mϕ/M-MDSCs. (A) Ly6G* neutrophil/PMN-MDSCs, p = 0.98. (B) Mo/Mϕ/M-MDSCs populations (Ly6C alternative M2, p = 0.33; Ly6C^int^ transitional, p = 0.70; and Ly6C^hi^ classical Ml, p = 0.63). (C) CCR2 MFI within Mo/Mϕ/M-MDSCs, p = 0.65. Data are mean ± SEM. NS = non-stressed, S = stressed; GD = gestational day; Mo/Mϕ/M-MDSCs = monocytes/macrophages/monocytic myeloid-derived suppressor cells; MFI = median fluorescence intensity.

**Supp. Fig. 2.**
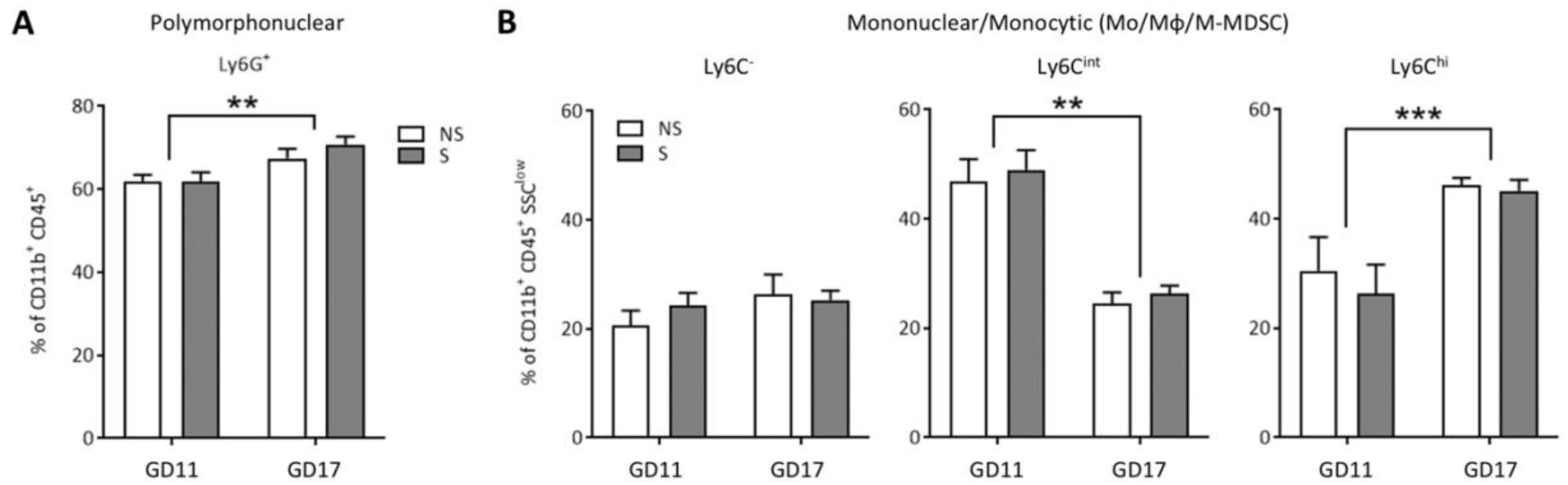
Placental polymorphonuclear and mononuclear populations differed across gestational time points (GD11 vs. GD17). Overall percent of (A) placental CDllb*CD45* polymorphonuclear (neutrophils or PMN-MDSCs) cells increased as gestation progressed from GD11 to GD17, regardless of prenatal stress (2-way ANOVA, main effect of gestational day, p = 0.0015). Overall percent of (B) SSC^low^ mononuclear cells (Mo/Mϕ/M-MDSCs) shifted towards an Ml pro-inflammatory Ly6C^hi^CCR2^hi^ phenotype (2-way ANOVA, main effect of gestational day, p = 0.0006), resulting in a decrease in the Ly6C^int^CCR2^int^ transitional phenotype (2-way ANOVA, main effect of gestational day, p < 0.0001) and no change in alternative M2 Ly6C CCR2 phenotype (2-way ANOVA, main effect of embryonic day, p = 0.19). Data are mean ± SEM. NS = non-stress, S = stress, GD = gestational day, Mo/Mϕ/M-MDSCs ≡ monocytes/macrophages/monocytic myeloid-derived suppressor cells. ** = p < 0.01, *** = p < 0.001.

**Supplemental Table 1.** Body weight and litter characteristics.

| Treatment | Non-stressed Control | N | Stress | N | p-value |
| --- | --- | --- | --- | --- | --- |
| Starting weight (g) | 21.7 ± 0.27 | 30 | 21.8 ± 0.27 | 34 | 0.80 |
| GD10 weight (g) | 24.9 ± 0.31 | 30 | 25.0 ± 0.27 | 34 | 0.73 |
| BW gained (g; GD0 □ GD10) | 3.2 ± 0.17 | 30 | 3.2 ± 0.18 | 34 | 0.86 |
| Litter size GD11 | 9.6 ± 0.32 | 8 | 9.3 ± 0.29 | 7 | 0.45 |
| Fetus resorption # | 0.13 ± 0.13 | 8 | 0 ± 0 | 7 | 0.35 |
| GD16 weight (g) | 32.6 ± 0.74 | 22 | 31.8 ± 0.40 | 27 | 0.36 |
| BW gained (g; GD0 □ GD16) | 11.1 ± 0.60 | 22 | 10.2 ± 0.41 | 27 | 0.18 |
| Litter size GD17 | 7.6 ± 0.46 | 22 | 8.1 ± 0.30 | 27 | 0.38 |
| Fetus resorption # | 0.59 ± 0.17 | 22 | 0.52 ± 0.12 | 27 | 0.73 |

Body mass, weight gain, and litter characteristics of non-stressed and stressed gestating dams. Unpaired T tests, using Welch's correction for unequal variances. Data are presented as mean ± SEM.

**Supplemental Table 2.** Spleen gene expression.

| Gene | Non-stressed | Stressed | p-value |
| --- | --- | --- | --- |
| *IL1B* | 1.0 ± 0.23 | 0.95 ± 0.15 | 0.85 |
| *IL6* | 1.0 ± 0.13 | 0.85 ± 0.11 | 0.43 |
| *IL10* | 1.0 ± 0.13 | 0.93 ± 0.14 | 0.76 |
| *IL23A* | 1.0 ± 0.20 | 0.90 ± 0.18 | 0.74 |
| *TNF* | 1.0 ± 0.14 | 1.17 ± 0.14 | 0.45 |
| *CCL2* | 1.0 ± 0.12 | 0.88 ± 0.04 | 0.28 |
| *TLR2* | 1.0 ± 0.16 | 1.32 ± 0.18 | 0.29 |
| *TLR4* | 1.0 ± 0.14 | 0.84 ± 0.12 | 0.43 |

Relative expression of immune genes in spleens of non-stressed and stressed gestating dams at GD17. Unpaired T tests, using Welch's correction for unequal variances. Data are presented as mean ± SEM; n = 5-10/group.

**Supplemental Table 3.** Circulating cytokines.

| Cytokine (pg/mL) | Non-stressed | Stressed | p-value |
| --- | --- | --- | --- |
| IL-1β | 2.56 ± 0.56 | 1.86 ± 0.40 | 0.32 |
| **IL-5** | **6.69 ± 1.13** | **3.74 ± 0.57** | **0.02 \*** |
| IL-6 | 13.79 ± 2.10 | 15.10 ± 2.20 | 0.68 |
| KC/GRO | 98.72 ± 9.46 | 91.91 ± 3.71 | 0.51 |
| IL-10 | 6.93 ± 0.45 | 6.29 ± 0.56 | 0.40 |
| TNFα | 7.64 ± 0.37 | 7.70 ± 0.35 | 0.90 |

Circulating cytokine levels (pg/mL) in the serum of non-stressed and stressed gestating dams at GD17, measured using MSD® multiplex mouse Proinflammatory Panel 1 assay. Restraint stress decreased circulating levels of Th2/mast cell cytokine IL-5 (an eosinophil and B-cell activator). Unpaired T tests, using Welch's correction for unequal variances. Data are presented as mean ± SEM; n = 12-16/group.

**Supplemental Table 4.** Uterine gene expression.

| Gene | Non-stressed | Stressed | p-value |
| --- | --- | --- | --- |
| *IL1B* | 1.0 ± 0.21 | 0.89 ± 0.14 | 0.66 |
| *IL6* | 1.0 ± 0.09 | 1.08 ± 0.12 | 0.65 |
| *IL10* | 0.81 ± 0.07 | 0.85 ± 0.08 | 0.68 |
| *IL2* | 1.0 ± 0.22 | 0.85 ± 0.21 | 0.64 |
| *CCL2* | 1.0 ± 0.19 | 1.02 ± 0.11 | 0.94 |
| *ITGAM* | 1.0 ± 0.13 | 0.82 ± 0.11 | 0.31 |
| *TLR4* | 1.0 ± 0.07 | 1.02 ± 0.05 | 0.77 |
| *TLR2* | 1.0 ± 0.08 | 0.87 ± 0.11 | 0.52 |

Relative expression of immune genes in uterine tissue of non-stressed and stressed gestating dams at GD17. Unpaired T tests, using Welch's correction for unequal variances. Data are presented as mean ± SEM; n = 6-10/group.

**Supplemental Table 5.**
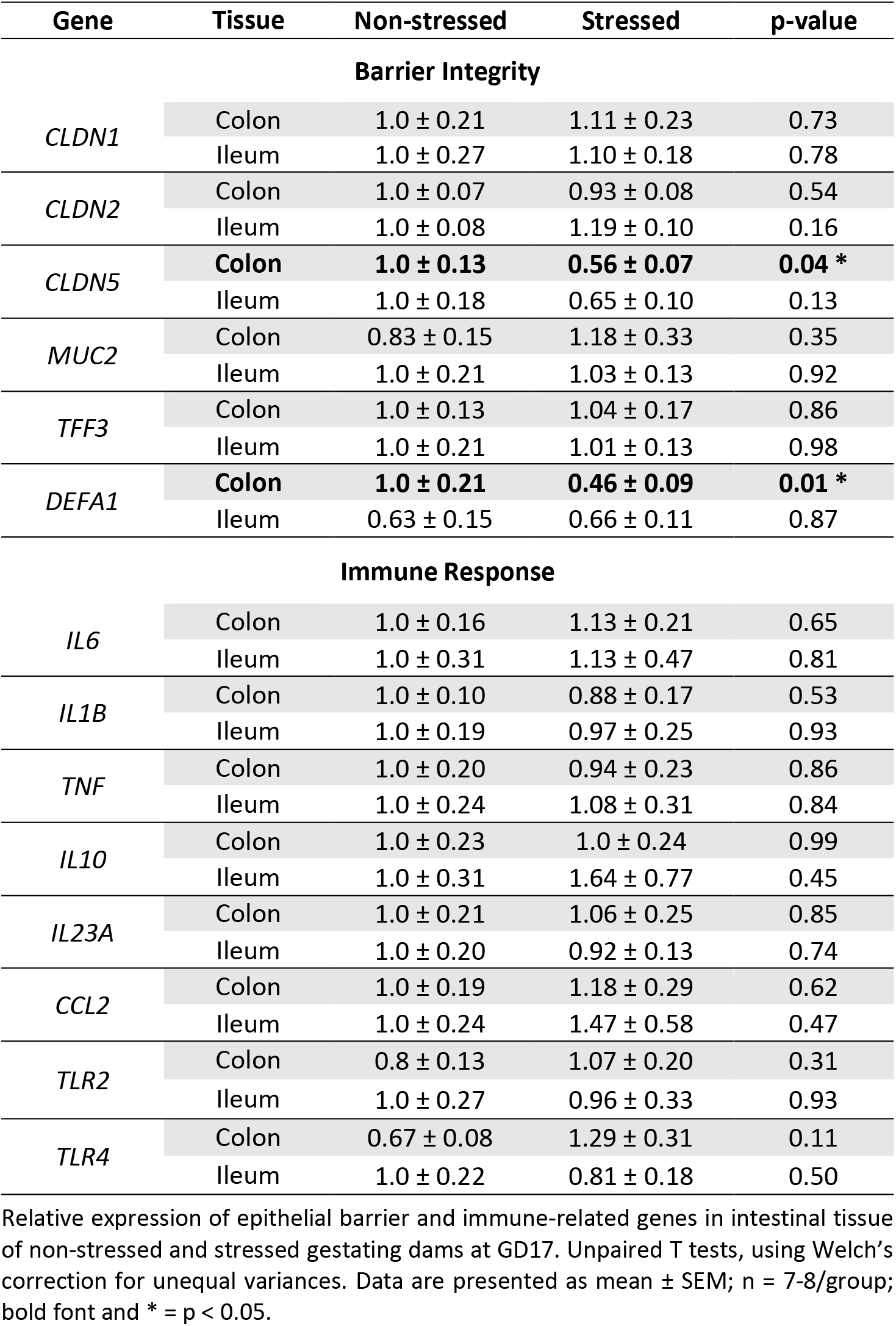
Intestinal gene expression.

**Supplemental Table 6.** Gene Primers.

| **Gene** | **Assay ID<sup>a</sup>** |
| --- | --- |
| *RPL19* | Mm02601633_g1 |
| *CRH* | Mm01293920_s1 |
| *NR3C1* | Mm00433832_m1 |
| *IL1B* | Mm00434228_m1 |
| *IL2* | Mm00434256_m1 |
| *IL6* | Mm00446190_m1 |
| *IL10* | Mm01288386_m1 |
| *IL23A* | Mm00518984_m1 |
| *ITGAM* | Mm00434455_m1 |
| *TNF* | Mm00443258_m1 |
| *CCL2* | Mm00441242_m1 |
| *TLR2* | Mm01213946_g1 |
| *TLR4* | Mm00445273_m1 |
| *CLDN1* | Mm00516701_m1 |
| *CLDN2* | Mm00516703_s1 |
| *CLDN5* | Mm00727012_s1 |
| *MUC2* | Mm01276696_m1 |
| *TFF3* | Mm00495590_m1 |
| *DEFA1* | Mm02524428_g1 |

<sup>a</sup> Applied Biosystems TaqMan Gene Expression Assay identification number.

